# Molecular signature of primate astrocytes reveals pathways and regulatory changes contributing to the human brain evolution

**DOI:** 10.1101/2023.12.12.570426

**Authors:** K. Ciuba, A. Piotrowska, D. Chaudhury, B. Dehingia, E. Duński, R. Behr, K. Soroczyńska, M. Czystowska-Kuźmicz, M. Abbas, I. Figiel, J. Włodarczyk, A. Verkhratsky, M. Niedbała, W. Kaspera, B. Wilczyński, A. Pękowska

**Author notes:** These authors contributed equally (K. Ciuba, A. Piotrowska). Correspondence to Aleksandra Pękowska.

## Abstract

Astrocytes contribute to the development and regulation of the higher-level functions of the brain, the critical targets of evolution. However, the molecular signature of foetal astrocyte evolution in primates is unknown. Here, to address this question, we use human, chimpanzee, and macaque induced pluripotent stem cell-derived foetal astrocytes (iAstrocytes). Human iAstrocytes are bigger and more complex than the non-human primate iAstrocytes. We find loci related to the regulation of cell size with increased expression in the human lineage. Likewise, we uncover that genes and mechanisms implicated in long-range intercellular signalling are activated in the human iAstrocytes. Strikingly, loci downregulated in the human lineage frequently relate to intellectual disability raising new questions on the trade-offs associated with the evolution of the human mind. Using our system, through a multilevel regulome analysis and machine learning, we uncover that functional activation of enhancers coincides with a previously unappreciated, pervasive gain of binding sites of ‘stripe’ transcription factors. In summary, we shed new light on a mechanism driving the acquisition of the regulatory potential of enhancers.

## Main

Astrocytes play essential and versatile roles in the central nervous system^1,2^ – they support the blood-brain barrier and maintain the proper composition of the extracellular milieu by, for instance, keeping the optimal concentration of ions and nutrients in the brain parenchyma ^1^. Astrocytes sustain the physiological level of neurotransmitters by their clearance, catabolism, and supply of obligatory precursors. In addition to these, broadly speaking housekeeping functions, astrocytes impact synaptogenesis^3–6^, synaptic activity^7,8^, and synaptic elimination ^9^ thereby contributing to the regulation of higher-level brain functions^10^.

Astrocytes have changed extensively in animal evolution ^11^ and there is a marked gain in astrocyte size and complexity in primates compared to rodents^11,12^. Single-cell transcriptomic analyses of post-mortem adult human and non-human primate (NHP) brains revealed that, compared to neurons, astrocytes feature more significant remodeling of the transcriptome^13–15^. Yet, a mechanistic and functional understanding of this phenomenon is largely missing.

Transcriptional changes linked to brain evolution often affect genes highly expressed during foetal and early postnatal development^16^. Notably, genes related to neurological disorders are also highly expressed during the embryonic period of the human’s life^17–19^. Yet, the access to foetal brain of our closest living relative the chimpanzee is impossible. Here, to address the molecular signature of foetal astrocyte evolution, we took advantage of the induced pluripotent (iPS) stem cells which yield foetal astrocytes robustly. We find that the human iPS-derived astrocytes (iAstrocytes) are larger and more complex than the NHP iAstrocytes. Using multiomics and machine learning, we map transcriptomic, epigenomic, and genetic signatures of the gradual and consistent mechanisms driving the evolution of astrocytes. Our data put astrocytes in the context of brain evolution and reveal grand axes shaping the alterations of functions of astrocytes in primates.

## Results

To obtain iAstrocytes, we opted for the monolayer-based approach as it allows to generate a substantial number of cells amenable to extensive functional tests. We took advantage of the well-established procedure to obtain cortical neural progenitors (iNP) and subsequently astrocytes (iAstrocytes)^20^ from a panel of primate iPS lines^21^ (**Fig. 1a, Ext. Fig. 1a-c, Methods**). The iAstrocytes expressed canonical astrocyte markers (**Fig. 1b**), including calcium-binding protein S100B, glial fibrillary acidic protein (GFAP) and Excitatory Amino Acid Transporter/glutamate transporter Solute Carrier Family 1 Member 3 (EAAT1/SLC1A3). When co-cultured with rat embryonic cortical neurons, all the iAstrocyte lines promoted synaptogenesis and neuronal survival of the rodent cells (**Ext. Fig. 1d-e**). Likewise, iAstrocytes took up glutamate from the extracellular environment (**Fig. 1d**) and responded to extracellular ATP by generating calcium (Ca^2+^) signals (**Ext. Fig. 1f**). To further assess the quality of our samples, we determined the expression of a comprehensive panel of astrocytic markers (GO:0048708) using bulk RNA sequencing (**Ext. Table 1**). We employed a well-established computational strategy^13,14,22^ and aligned the paired-end RNA-seq reads to a consensus genome built upon the most recent human, chimpanzee, and rhesus assemblies (**Methods**). All the primate iAstrocytes featured a pattern of expression of astrocytic genes closely resembling that of the human primary foetal astrocytes cultured in vitro and of cells previously obtained from a panel of distinct female iPS lines^20^ (**Fig. 1c**).

**Figure 1:**
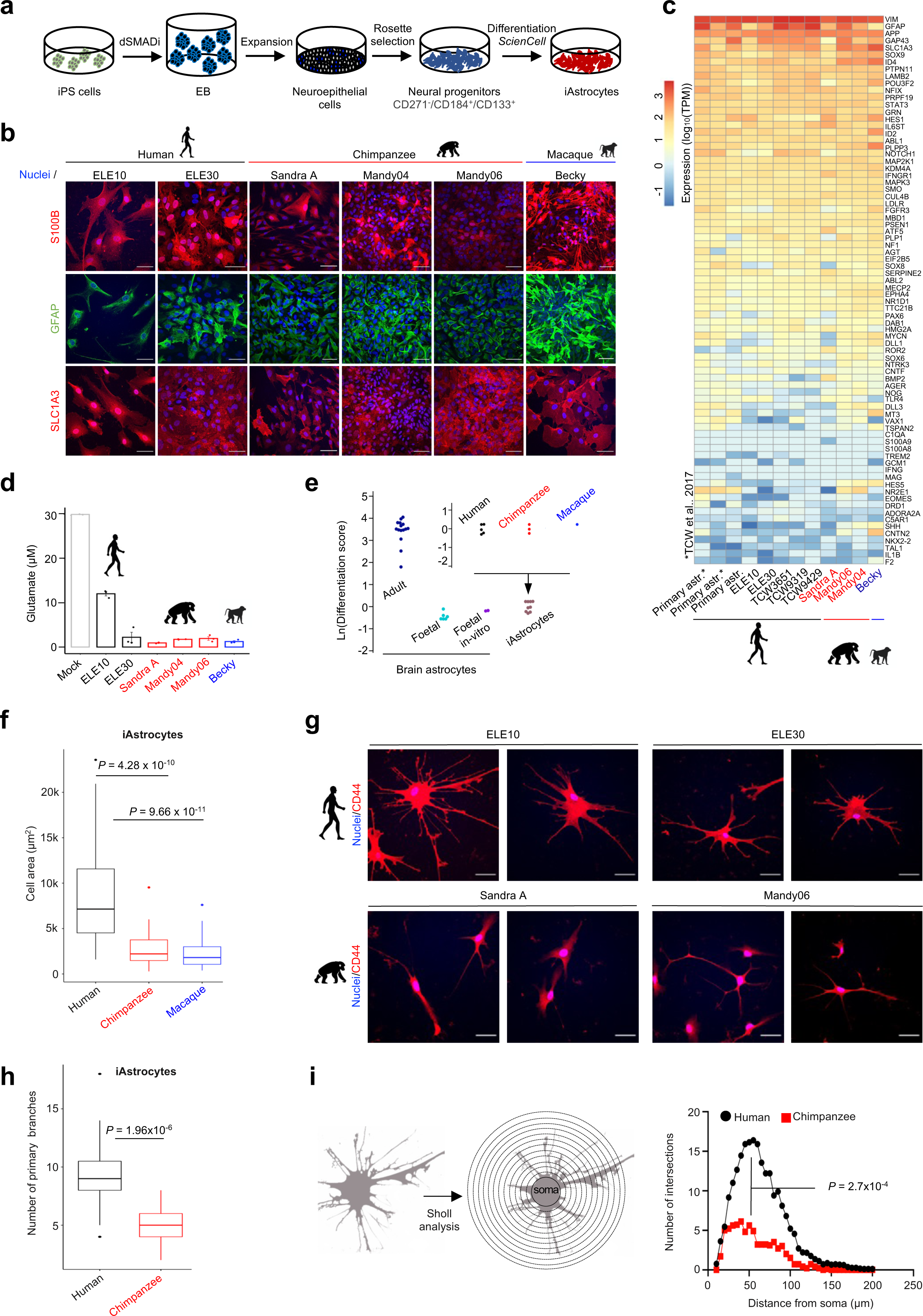
Primate iAstrocytes recapitulate evolutionary features of foetal brain astrocytes. **a.** Experimental strategy to obtain primate (human, chimpanzee, macaque) iAstrocytes. **b.** Representative immunofluorescence images of primate iAstrocytes showing the expression of canonical astrocytic markers: S100B (red), GFAP (green), SLC1A3 (red). Scale bar: 50μm. **c.** Heatmap displaying the RNA-seq (TPM, transcript per million) expression level of a cohort of astrocyte markers in human foetal brain-derived astrocytes cultured in vitro and in primate iAstrocytes. **d.** Glutamate uptake from the extracellular milieu by iAstrocytes (n=3). **e.** Comparison of differentiation score (DS) of primate iAstrocytes reveals similar degree of maturation of cells across lines and species. Through quantitative comparison of the published RNA-seq data for foetal and adult brain-derived astrocytes^28^, we identified genes overexpressed in immature and mature astrocytes (foetal and adult markers respectively, *P*-adj. < 0.01, log2(Adult/foetal) | > 1, *DESeq2* method). We defined DS as logarithm base 2 of the ratio between the average expression level of adult and foetal markers. The dot-plot displays DS for samples grouped by experiment and species. **f.** Quantification of the total cell area of tdTomato-expressing iAstrocytes (n=43 (human), 51 (chimpanzee), 47 (macaque); *P*-value: two-sided t-test). **g.** Human iAstrocytes feature more complex morphology than their chimpanzee counterparts. Representative examples of CD44-immunostained human and chimpanzee iAstrocytes. Scale bar: 50μm. **h.** Number of primary projections of human and chimpanzee CD44-stained iAstrocytes (n=28 (human); 19 (chimpanzee); *P*-value: two-sided t-test). **i.** Left panel: scheme for Sholl analysis of iAstrocytes. Intersections were measured every 5 μm in 200 μm radius, starting from the cell soma (concentric circles); right panel: average number of intersections in the function of the distance from the cell soma; n=20 (human), 14 (chimpanzee); *P*-value: two-sided t-test).

To capture the maturation status of the of iAstrocytes, we took advantage of published transcriptomes of acutely purified foetal and adult human brain astrocytes^23^ and defined *differentiation score* (DS; **Methods**). We found that iAstrocytes from the three species displayed a DS indicating a foetal astrocyte identity. Critically, all the preparations featured a similar DS testifying a comparable ‘molecular age’ of human, chimpanzee, and macaque iAstrocytes (**Fig. 1e**). Altogether, these data testified successful conversion of the iNP cells to functional foetal astrocytes.

### iAstrocytes recapitulate the salient features of astrocyte evolution

In the adult brain, human astrocytes feature more complex morphology in comparison to rat, mouse, and macaque ^11,12^. We found that the human iAstrocytes had a significantly larger cell surface area than chimpanzee and macaque cells (**Fig. 1f-g, Ext. Fig. 1g**, two-sided t-test, Human vs Chimpanzee *P* = 4.28 x 10^-10^; Human vs Macaque *P* = 9.66 x 10^-11^). In contrast, we found no significant difference in the surface area of nestin-positive iNP cells of the corresponding species (**Ext. Fig. 1h-i**). Remarkably, human iAstrocytes were endowed with, on average 1.74 times more primary branches than the chimpanzee iAstrocytes (9.3 versus 5.3, two-sided t-test *P*=1.96×10^-6^, **Fig. 1g-h**). Sholl analysis, which measures the number of intersections of cellular projections at different distances from the cell soma, revealed that human iAstrocytes exhibited on average 3 times more intersections of processes than their chimpanzee (5.64 versus 1.88, *P* = 2.7×10^-4^, two-sided t-test, **Fig. 1i**). Thus, the in vitro system recapitulates the interspecies differences in size and complexity of astrocytes thereby capturing the salient features of astrocyte evolution. Likewise, our data reveal the genetic encoding of the enhanced astrocyte complexity in evolution.

### Changes in gene expression in astrocyte evolution

Differential gene expression is the major contributor to the evolution of traits^24,25^. As we established that the in vitro system allows studying the evolution of astrocytes and that the ‘molecular age’ of our samples is similar across all the species, we next sought to define genes, the expression of which distinguishes human from NHP iAstrocytes. To increase the robustness of our approach, we considered both our and the published RNA-seq data of three additional human female iAstrocyte lines^20^ obtained using the same protocol (**Fig. 1c**).

Multiple lines of evidence testified our capacity to capture evolutionary changes in astrocyte transcriptomes: (*I*) consistent with the evolutionary distance separating species included in our study, there were significantly more differentially expressed genes (DEG) in the human-to-macaque than in the human-to-chimpanzee iAstrocyte comparison (**Fig. 2a**, **Ext. Table 2**); (*II*) we scored a largely congruent transcriptional changes for genes affected in both comparisons (*P*-adj. < 0.01, | logarithm base 2 of fold change (LFC) | > 0, n=1,271, Pearson’s *r*=0.77, *P*<2.2×10^-16^, **Ext. Fig. 2a**), which is in agreement with the previously published reports^13,14^; (*III*) an overwhelming majority of the EAGs were robustly expressed in iAstrocytes and in acutely purified brain astrocytes^23^ (TPM>1, **Fig. 2b**, **Ext. Fig. 2b, Methods**); (*IV*) as expected ^14^, we observed transcriptional modulation rather than dramatic changes in gene expression in evolution (**Ext. Fig. 2b**) which is consistent with a stronger evolutionary constraint on the protein coding gene expression. We identified 493 up- and 373 downregulated EAGs, while comparing human with NHP iAstrocytes (**Fig. 2c**). Altogether, these data are largely consistent with predictions and highlight our power to uncover evolutionary changes in astrocyte biology.

**Figure 2:**
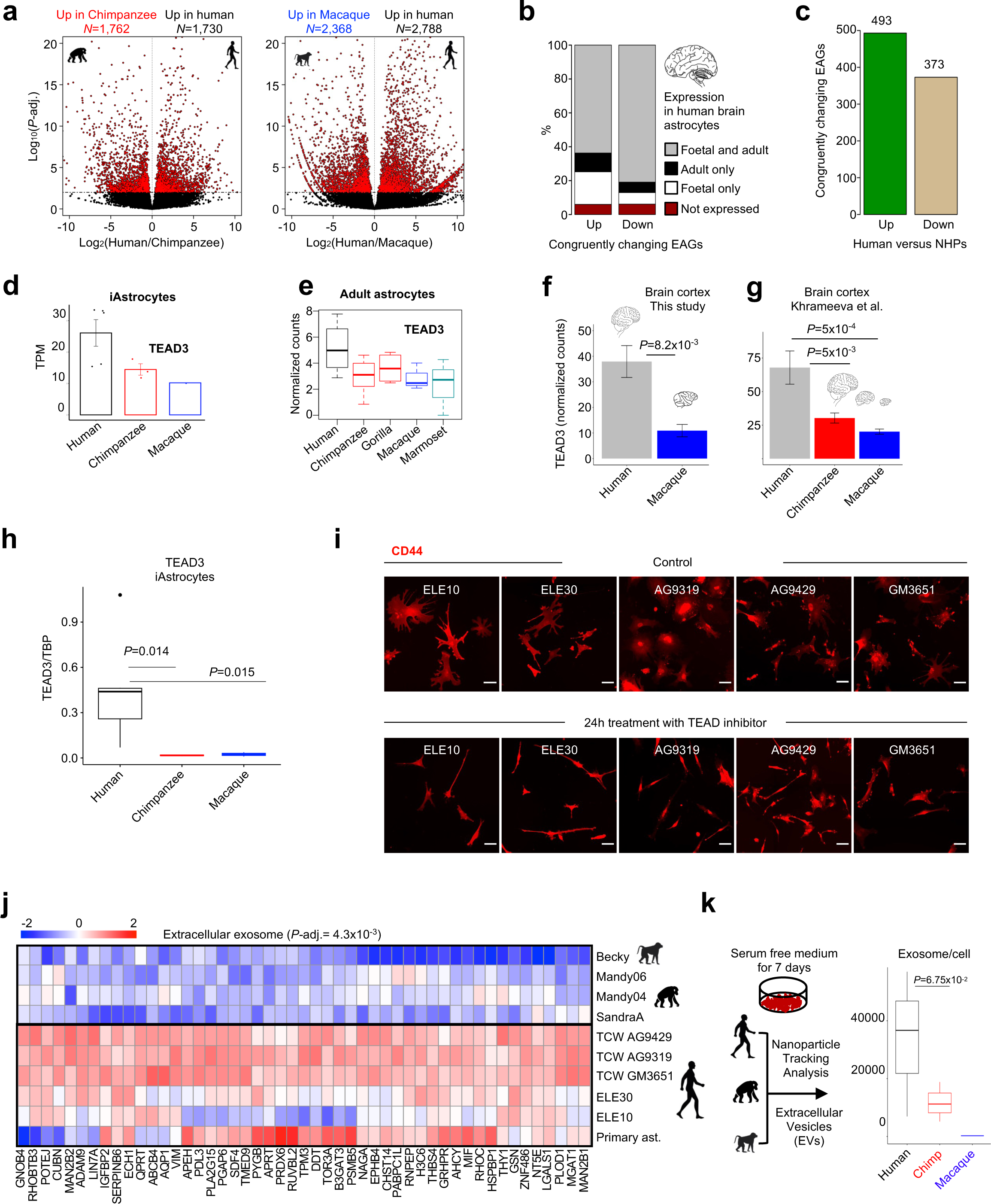
Stepwise evolutionary changes in gene expression during astrocyte evolution. **a.** Transcriptional differences between human and NHP iAstrocytes. Volcano plot illustrating the results of the *DESeq2-*based identification of differentially expressed genes (DEG) in human versus chimpanzee and in human versus macaque iAstrocytes (red denotes adjusted *P*-val.<0.01). We considered DEG that change expression in a congruent manner in the two comparisons and focused on loci that feature high and consistent level of expression in iAstrocyte lines (upregulated in humans: TPM>1 in at least 4 out of 5 human iAstrocyte lines; downregulated in humans: TPM>1 in at least 3 out of 4 NHP samples (n_up_ = 677; n_down_=486). **b.** Expression of loci identified in panel **a** in acutely purified foetal and adult brain astrocytes^23^. 612 out of the 677 upregulated (90%) and 448 out of the 486 downregulated (92%) EAGs were robustly expressed in acutely purified human brain astrocytes. **c.** Final numbers of EAGs identified in human vs NHP iAstrocytes considered in the functional analyses. To make sure any possible residual differences in maturation did not impact subsequent functional analyses, foetal astrocyte markers were removed from the list of upregulated EAGs, while adult astrocyte markers were removed from the list of downregulated EAGs. **d.** Expression (TPM) of TEAD3 in primate iAstrocytes. *TEAD3* is an EAGs linked with the Hippo pathway. Hippo pathway regulates organ size. **e.** Expression (normalized RNA-seq counts, *DESeq2* method) of TEAD3 in human, chimpanzee, gorilla, macaque and marmoset adult cortical brain astrocytes from the middle temporal gyrus (data from ref 15). **f.** Expression (normalized RNA-seq counts, *DESeq2* method) of TEAD3 in human and macaque brain cortical tissue samples. **g.** Expression (normalized RNA-seq counts, *DESeq2* method) of TEAD3 in human, chimpanzee, and macaque brain cortical tissue samples (published data set, included are only female samples). **h.** Protein level of TEAD3 is increased in the human as compared to chimpanzee and macaque iAstrocytes (*P,* two-sided t-test). **i.** Inhibition of TEAD leads to diminished cell size and complexity of iAstrocytes. Five iAstrocytes lines were treated for 24h with 3 μM VP-107, a pan TEAD inhibitor. Cells were then fixed and immunolabelled for the expression of CD44 (red, scale bar – 50μm). **j.** Heatmap of scaled expression of EAGs linked to extracellular exosome, including foetal brain astrocytes. **k.** Number of exosome per cell in primate iAstrocytes quantified with NTA (n=3, *P*-value: two-sided t-test).

### *TEAD3* expression is evolutionarily gained in the human iAstrocytes and in the human brain

It is currently unknown which genes regulate the increase in astrocyte size and complexity in evolution. We found several examples of genes related to the regulation of cytoskeleton (**Ext. Fig. 2d**). Notably, our results revealed that the expression of TEA domain transcription factor 3 (*TEAD3*), one of the direct targets of the Hippo pathway that controls cell and organ size^26,27^, is evolutionarily gained in the human iAstrocytes (**Fig. 2d**). Yet, whether the Hippo pathway might be related to the evolutionary changes in cell size remains unknown.

Recent single-cell RNA-seq profiling of astrocytes in the middle temporal gyrus in five primate species^15^ confirmed transcriptional activation of *TEAD3* in the human adult brain astrocytes compared to astrocytes from all four non-human primate samples assessed in that study (**Fig2. e).** Thus, the transcriptional upregulation of *TEAD3* in the human lineage is maintained in the adult brain astrocytes. Moreover, we obtained transcriptome profiles from adult human and macaque brain cortices and determined the expression change of *TEAD3*. Human samples featured a significantly higher activity of *TEAD3* gene (**Fig. 2f**, LFC=1.8, *P*-adj.=8.2×10^-3^, *DESeq2* method). We then compared published transcriptomes from human, chimpanzee, and macaque female brains (**Methods**). *TEAD3* was expressed at the highest level in the human samples (**Fig. 2g**, LFC human vs. chimpanzee=1.16, *P*-adj.=5×10^-3^, *DESeq2* method). We then assessed the protein level of TEAD3 in iAstrocytes by Western blot and confirmed its increased in human iAstrocytes (**Fig.2h**).

Next, we determined the contribution of the activity of TEAD proteins in the regulation of astrocyte size and complexity. We obtained and validated iAstrocytes from additional human female iPS lines ^28^ (not shown) and incubated iAstrocyte preparations for 24 hours with VT-107, a selective inhibitor of TEAD proteins (3µM). Strikingly, while having no impact on cell survival in all the iAstrocyte lines, inhibition of TEADs led to significantly decreased cell size and diminished cell complexity featuring a drastically reduced number of branches in all the assayed iAstrocyte lines (**Fig. 2i**). Altogether, through our analysis, we identified TEAD activity as a contributor to astrocyte size and complexity. We hypothesize that the transcriptional upregulation of *TEAD3* may help expand astrocyte size and the number of astrocyte processes in evolution.

### Evolutionary gain of expression of genes related to extracellular vesicles

Bulk and single-cell analyses of gene ontologies (GO) of loci activated in the adult human as compared to the NHP brain revealed frequent alterations in expression levels of genes related to synapse biology^13,29^, and, in mature astrocytes, in genes regulating synapse production and activity^30^. However, it remains unclear which cellular pathways are enhanced in foetal astrocyte in the course of evolution. To address this question, we considered a broad GO database (https://david.ncifcrf.gov) and assessed the enrichment of GO terms and signaling pathways in the lists of up- and downregulated EAGs.

We found a significant overrepresentation of genes related to metabolism amongst the upregulated EAGs (**Ext. Fig. 2c**) including mitochondrial ribosomal protein S7 (*MRPS7*), mitochondrially encoded ATP-synthases 6 and 8 and factors implicated in lipid signaling (phospholipase C delta 3 – *PLCD3*, phospholipase D family member 3 – *PLD3* and phospholipid scramblase 3 – PLSCR3, **Ext. Fig. 2c**). This result is in line with the fact that the evolution of the primate brain is marked by an increase in metabolic expenditure ^31,32^. Human and the NHP brain consume ∼20%, and ∼10% of the whole body’s metabolic resources, respectively ^33–35^. Thus, foetal astrocyte evolution is related to gain in expression of genes regulating the production and consumption of energy.

Strikingly, 49 out of 493 (10%) of upregulated EAGs were related to the extracellular exosome ontology (ExEg, *P-*adj.= 4.3×10^-3^, **Fig. 2j**) including, for instance, EPH receptor B4 (*EPHB4*), a positive regulator of neuron projection and Thrombospondin 4 (*THBS4*). *EPHB4* has recently been shown to be activated in the human as compared to NHP adult Middle Temporal Gyrus astrocytes^30^. Thrombospondins are secreted glycoproteins regulating synaptogenesis and THBS4 is a relatively poorly understood member of this family. Previous reports revealed increased expression of THBS4 protein in human as compared to chimpanzee, and macaque brains^36^, which corroborates our results.

Genes in the ExEg category included loci implicated in the biology of exosomes – the extracellular structures that impact homeostasis and pathology of the nervous system^37–39^. To determine if exosome production differs between iAstrocytes from the three primate species, we isolated exosomes from the culture media of human and NHP iAstrocytes and quantified them using Nanoparticle Tracking Analysis (NTA, **Methods**). The production of exosomes was higher in chimpanzee as compared to macaque and was further enhanced in the human iAstrocytes (**Fig. 2k**). Thus, the evolution of primate astrocytes frequently triggers genes related to the extracellular milieu and manifests itself with enhanced production of exosomes. Changes in the composition of the extracellular milieu are likely an integral facet of brain evolution, with likely implications in synaptogenesis.

### Downregulation of genes related to chromatin biology in primate astrocyte evolution

The analysis of ontologies linked to downregulated genes in the human iAstrocytes, revealed that 249 out of 373 downregulated EAGs (67%) were nuclear factors (DAVID/KEGG database; Fold enrichment=1.9, B-H adjusted *P*=1.4×10^-17^, **Ext. Fig. 3a**). This included sequence-specific transcription factors such as zinc finger KRAB regulators and chromatin remodellers. We observed a significant enrichment of genes encoding negative regulators of splicing (Fold enrichment=17.9, B-H adjusted *P*-adj.=6.6×10^-3^). In agreement with the published analyses in the adult animal brain tissue^40^, we found a stepwise increase in the number of splicing events in the human as compared to chimpanzee and macaque cells (**Ext. Fig. 3b, Methods**). Taken together, evolution frequently leads to diminished expression of nuclear factors including genes that negatively impact splicing which might contribute to the observed increased complexity of the mRNAs produced in the human iAstrocytes.

### Genes related to intellectual disability / mental retardation are silenced during evolution

The propensity to succumb to specific neurological disorders might arise corollary of brain evolution^41,42^. Thus, we sought to determine how evolutionary changes in gene expression in astrocytes might be related to human diseases. We queried the most recent, manually curated database of 5,669 genes related to brain malfunction (BrainBase; https://ngdc.cncb.ac.cn/brainbase/index). EAGs were related to a total of 23 diseases including Glioma, Alzheimer’s disease, Autism Spectrum Disorder, or Multiple Sclerosis (**Fig. 3a**). While the total number of observed disease-related EAGs (n*_observed_*=131) matched the expected value (n*_expected_*=113, **Fig. 3a, inset**), we found a 2.5-fold overrepresentation of downregulated genes amongst the disease-related EAGs (*P*=3.4×10^-6^, Fishers’ exact test, 85 down and 46 upregulated EAGs).

**Figure 3:**
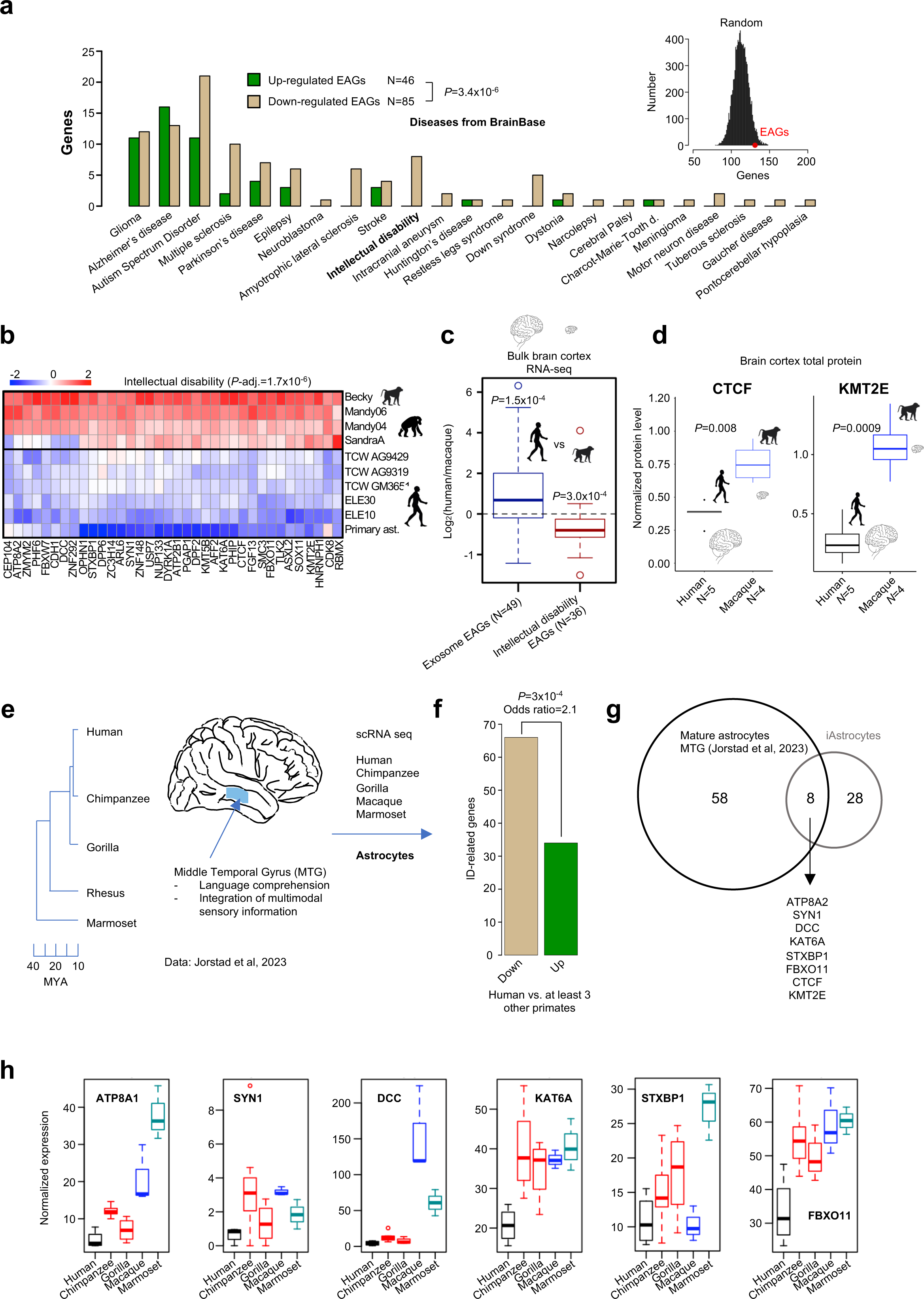
Intellectual disability genes are frequently transcriptionally downregulated in the evolution of the human brain. **a.** EAGs linked to human brain disorders. Downregulated EAGs are more frequently linked to brain disorders than the activated EAG (Chi^2^ test *P*-val.=5×10^-8^). Inset – number of EAGs related to diseases matches the expected value. Displayed is the distribution of the number of genes related to brain disorders in sets of 866 genes (493+373) sampled randomly from the genome. **b.** Heatmap of scaled expression of downregulated EAGs related to ID in iAstrocytes and in foetal astrocytes (ID genes from the UniProt database, Benjamini Hochberg corrected *P*-value was obtained through https://david.ncifcrf.gov/). **c.** Exosome-linked and ID-related genes identified in **Fig.2h** and **Fig.3b**, respectively feature a pervasive transcriptional upregulation in the human (n=6) as compared to macaque brain cortex (n=2). Inverse is true for ID-related genes. Bulk RNA-seq was obtained from human and macaque cortical tissue. **d.** Protein level of ID-related CTCF and KMT2E (left and right panels respectively) is diminished in the human as compared to macaque brain cortex tissue (*P* two-sided t-test). **e.** Analysis single cell RNA-seq obtained for primary post-mortem Middle Temporal Gyrus of human, chimpanzee, gorilla, rhesus macaque, and marmoset cortices (data from Jorstad et al., 2023). Cells annotated as astrocytes were considered. Differentially expressed genes were retrieved (*P*-adj<0.1, arrow). **f.** ID-related genes are more frequently transcriptionally down regulated than up-regulated in human as compared to non-human primate MTG astrocytes. Displayed is the P-value from Fisher’s exact test. **g.** Venn diagram depicting the overlap between evolutionarily downregulated genes in the human mature astrocytes in the MTA (analysis as in panel f) and human iAstrocytes. **h.** Expression of a panel of genes concordantly downregulated in the human astrocytes and iAstrocytes in primate MTG astrocytes (n=5 for human; n=7 for chimpanzee; n=4 for gorilla; n=3 for rhesus; and n=3 for marmoset); data from Jorstad et al., 2023.

Remarkably, all the EAGs related to intellectual disability (ID) from the BrainBase were within the transcriptionally downmodulated group (*P*=4.6×10^-3^, Pearson’s χ^2^ test, **Fig. 3a**). We performed additional analyses and experiments to better define the link between ID and evolutionary changes in astrocyte transcriptome. We considered the broadest annotation of genes related to ID (https://www.uniprot.org/keywords/KW-0991). While only 1 ID-related gene in the Uniprot database was activated in the human cells, 36 ID-related loci were amongst the EAGs downregulated in the human iAstrocytes (odds=47, *P*=3.7×10^-12^, Fishers’ exact test, **Fig. 3b**). To test whether the link between ID and evolutionary alterations in gene expression are analogous in the adult brain samples, we considered our and published ^14^ bulk RNA-seq libraries from human, chimpanzee, and macaque frontal cortices. The 36 ID-related genes featured overall downregulation in the adult human brain as compared to the NHP samples while *loci* related to the extracellular exosome were by and large activated in human samples (*P*=1.5×10^-4^ and *P*=3×10^-4^ respectively, two-sided t-test **Fig. 3c, Ext. Fig. 3c**). Conforming to these analyses, western blot assessment of the cortical grey matter in human and macaque adult brains revealed a significant decrease in protein levels of two chosen ID-related genes CTCF and KMT2E in the human samples (**Fig. 3d**, **Ext. Fig. 3f**).

We next asked if the evolutionary downregulation of ID-related genes was also significant in the adult astrocytes from the middle temporal gyrus (MTG), a cortical region implicated in the integration of multisensory input and in the comprehension of language (**Fig. 3e**). In line with our findings, ID-related genes were 2-times more frequently down-regulated than activated (*P*-adj.<0.1 in at least three out of four inter-species comparisons) in the human MTG adult astrocytes (**Fig. 3f**; *P*=3×10^-4^, Fisher’s exact test). We found a significant overlap between the ID-related genes downregulated in the human adult astrocytes and in iAstrocytes (**Fig. 3g-h**). Thus, ID-related genes are frequently transcriptionally downregulated in human astrocyte evolution.

Next, we addressed the inverse question and determined the LFC of expression of all genes related to each of the neurological disorders in human compared to the NHPs brain cortex. ID constituted the only condition for which we observed a congruent and consistent significant downregulation of expression in all the comparisons (**Ext. Fig. 3d-e**, n*_genes_*=138). Taken together, evolution frequently leads to downregulation of the expression of genes critically important for higher-level brain function, and their dosage is the key to proper mental abilities of the human brain.

### Transcriptional changes in astrocyte scale with the net gains in the number of enhancer elements in a context of preserved local three-dimensional chromatin topology

Variations in transcriptional regulatory systems may drive the acquisition of new traits^24,25,43^ and the analysis of interspecies differences in chromatin accessibility helps reveal mechanisms orchestrating the alteration of gene expression in evolution^13,44–47^. We adopted an ATAC-seq-based approach to best account for the evolutionary changes in cis-regulatory element (CRE) activity. Likewise, we mapped H3K4me3 and H3K27ac histone modifications which are enriched at active promoters (H3K4me3, H3K27ac) and active enhancer elements (H3K27ac)^48,49^.

Validating our capacity to emulate the changes in chromatin activity across astrocyte evolution, the principal component analysis (PCA) revealed tight clustering of regulomes by species (**Fig. 4a**). Changes in openness at intergenic ATAC-seq regions between human and chimpanzee and human and macaque iAstrocytes were less correlated than the alterations in gene activity^50–52^ (**Ext. Fig. 4a-b**, Pearson’s *r*=0.42 for ATAC and *r*=0.77 for RNA).

**Figure 4:**
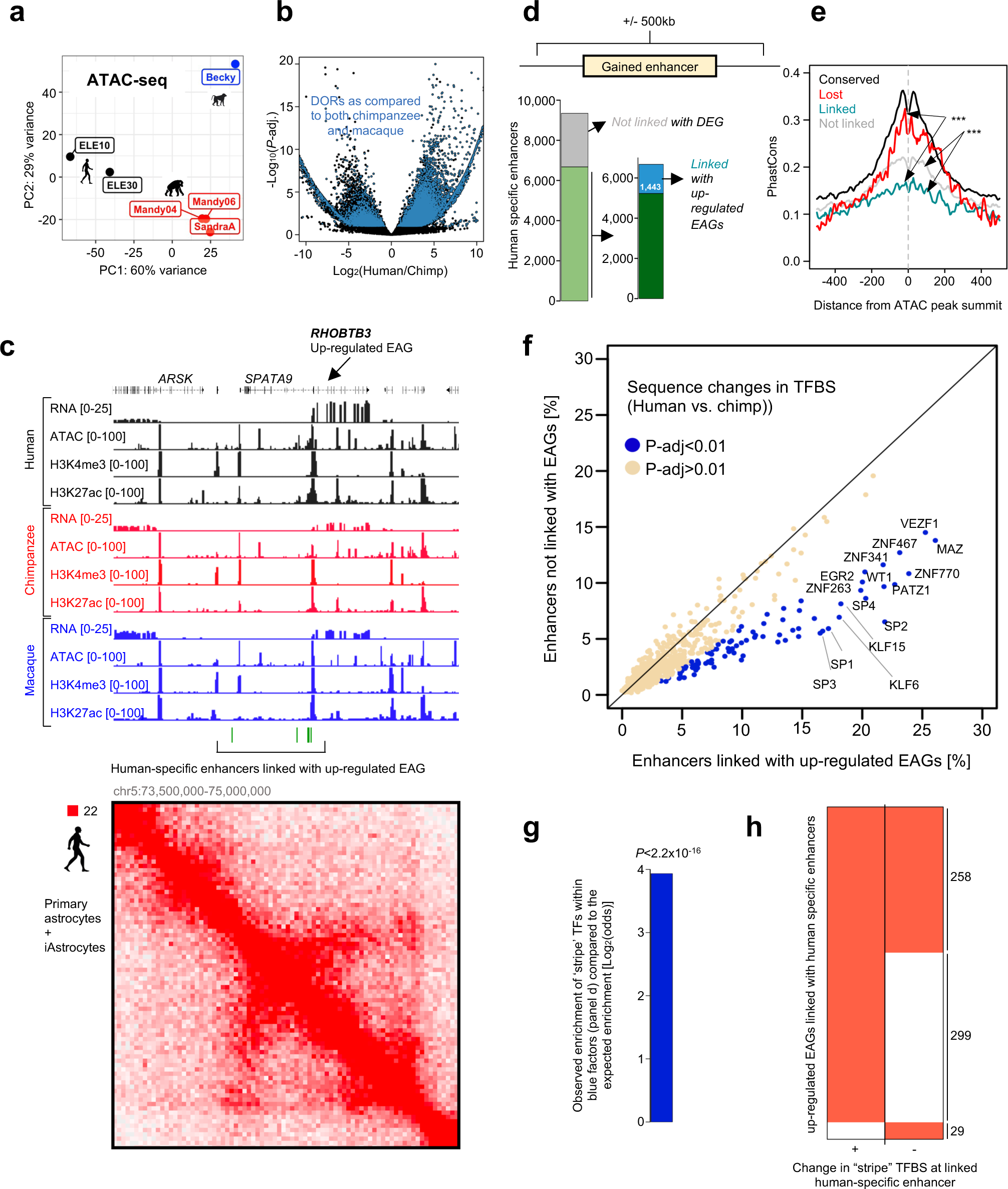
Gain of binding sites of ‘stripe’ transcription factors at enhancers linked to gene activation. **a.** Principal component analysis (PCA) reveals clustering of human, chimpanzee, and macaque iAstrocyte profiles of chromatin openness by species. Filtered ATAC-seq reads were counted within genomic intervals corresponding to peaks detected in at least one species which were also aligneable between the three genomes (minimum 50% of the peak interval was transferable between genomes; *N*=224,411 regions). PCA was performed after data normalisation (*DESeq2*). **b.** Volcano plot illustrating the analysis of differential openness in primate iAstrocytes. We identify regions that vary ATAC-seq signal between human and the two NHP iAstrocyte lines (blue indicates significant and congruent change in human as compared to chimpanzee and macaque cells, *P*-adj.<0.1, *DESeq2* method, for simplicity, only human-chimpanzee comparison of LFC and *P*-values is shown). **c.** Gene expression, along with chromatin activity and architecture, at the RHOBTB3 locus. RHOBTB3 is an upregulated EAG surrounded by human-gained enhancers (green bars); upper panel: RNA-, ATAC- and ChIP-seq tracks aligned to the consensus genome; lower panel: in-situ-Hi-C reads aligned to the human genome. **d.** Majority of enhancers gained in the human iAstrocytes can be linked with genes varying expression in evolution. Enhancers gained in the human iAstrocytes were stratified into groups based on the presence of promoters of differentially expressed genes (DEGs) in their genomic vicinity (+/-500 kb). The ‘linked’ enhancers were enhancers located within 500kb of an upregulated EAGs. The ‘not-linked’ enhancers were elements for which there was no promoter of DEG in the vicinity (+/-500 kb; we considered all the DEGs scored in the comparison between human and chimpanzee and in the comparison between human and macaque iAstrocytes to define the ‘non-linked’ elements). **e.** PhastCons score (average) in the genomic intervals surrounding the summit of the common, lost and gained ATAC peaks (*** P < 0,001, two-sided t-test). **f.** Per TF frequency of elements featuring changes in the predicted transcription factor binding sites (TFBS) differs between human-specific enhancers linked with upregulated EAGs and the human-gained elements not linked with differential gene expression in primate iAstrocytes. The DNA sequences of each enhancer were compared between human and chimpanzee. Discordant positions (mismatches, insertions, or deletions) intersecting predicted TFBS were identified. Plot compares, for each TFs and in each of the two groups of enhancers, the percent of elements that feature an evolutionary difference in sequence. Blue: TFs being significantly more frequently changed at enhancers gained in the human lineage and linked with upregulated EAGs than at elements not linked with DEGs in primate iAstrocytes (P-adj.<0.01; Fisher’s test; n=89). **g.** Enrichment of ‘stripe’ factors within the group of TFs identified in panel d (blue; P-value from Fisher’s test); 77 out of 89 TFs showing a significantly higher frequency of evolutionary changes within the enhancers linked to upregulated EAGs are ‘stripe‘ TFs. **h.** Overwhelming majority of upregulated EAGs are located within 500kb of an enhancer gained in the human lineage that features an evolutionary change in the motif of one of the 77 ‘stripe’ TFs identified in panel F.

We identified 16,496 differentially open regions (DORs, *P*-adj.<0.1 in both comparisons, *DESeq2* method, **Fig. 4b, Ext. Table 2**) and observed stepwise evolutionary changes in chromatin openness at DORs (**Ext. Fig. 4c-d**). As predicted, most DORs were located outside promoters which is consistent with the essential role of distal regulatory elements in eliciting evolutionary changes in gene expression^43–45^. We identified 9,343 human- and 2,351 NHP-specific enhancers (**Methods**).

Cognate promoter-enhancer pairs tend to reside in shared topologically associating domains (TADs), and it is proposed that TADs constitute functional units of genome organisation ^53–56^. Thus, to better define the possible mechanisms regulating gene expression in astrocyte evolution, we performed high-resolution *in-situ* Hi-C^57^. We obtained a total of 4.7 billion reads and mapped TADs in human primary (foetal) brain astrocytes and in human and chimpanzee iAstrocytes (**Ext. Fig. 4e, Ext. Table 3**). The annotations of TAD boundaries were vastly congruent between primary brain astrocytes and iAstrocyte samples (**Ext. Fig. 4e**) certifying the quality of the data and further highlighting our power to mirror regulomes of astrocytes. Consistent with the published results ^58–60^, TAD architecture was globally preserved in primate astrocytes (**Ext. Fig. 4e-f**) and de novo formation of TAD boundaries was rare (**Ext.Fig. 4f**). Rather, the species-specific boundaries (**Ext. Table 3, Methods**) arose seemingly as a consequence of an increase in the insulation strength of sites that featured boundary potential in the other species (**Ext. Fig. 4g**). We therefore focused on the interplay between evolutionarily dynamic regulome and transcriptomes taking into consideration TAD annotation in the human iAstrocytes.

Remarkably, TADs by and large contained genes changing expression in a congruent manner in evolution: out of 157 TADs that featured at least 2 EAGs, only 33 domains (21%) contained both up- and downregulated EAGs while 124 (79%) featured loci changing in the same direction (3.7-fold overrepresentation, **Ext. Fig. 4h**) suggesting a cooperative evolutionary gain in expression of genes in shared TADs.

Consistent with previously published results^13^, TADs featuring promoters of upregulated EAGs also frequently embedded human-specific enhancers (253 of 415; 61%, **Ext. Fig. 4i**), while an overwhelming majority of domains intersected an enhancer that varied openness in any of the two comparisons (358/415; 86%). The number of enhancers gained in each interspecies comparison correlated not only, as anticipated^61^, with the LFC of gene expression but also with the number of species-specific enhancers (**Ext. Fig. 4j-k**). Altogether, the dynamic reconfiguration of regulomes at loci changing expression in evolution entails stepwise gain of enhancer openness within otherwise largely preserved TADs. Our analysis reveals that the number of enhancers gained in the human lineage scales with the transcriptional onset and evolutionary distance between the species. Likewise, there is a congruent activation of gene expression in TADs in evolution.

### Gain of binding sites for ‘stripe’ transcription factors at enhancers is linked to gene activation

Changes in the sequence of transcription factor binding sites (TFBS) impact enhancer activity and shape gene expression in evolution^62,63^. By analogy, species-specific enhancers typically feature poorer conservation^50,61^. What is the nature of evolutionary-driven enhancer sequence changes that underlies activation of enhancers remains a fundamental question.

Previously, comparisons of neural crest cell regulomes, in human and chimpanzee, revealed targeting of unique TFBS at newly evolved enhancers^61^. We found that both the conserved elements and enhancers activated in the human iAstrocytes were enriched, amongst others, in motifs recognised by RUNX2, FOS/JUN, NFIA, NFIC transcriptional regulators known for their roles in astrocyte biology (not shown). However, whether there is a specific group of TFs able to endow the evolutionarily gained enhancers with a transactivatory potential is at present unknown.

To address this question, we sought of a conceptually new approach in which we search for sequence changes that contribute to genuine activation of enhancers, as evidenced by the changes in gene expression occurring in their genomic vicinity. In contrast to previous studies, as the expected background, we considered elements that were gained in the human lineage but featured seemingly no cis-regulatory activity.

Indeed, out of 9,343 ehancers activated in the human lineage only 1,443 (15%) could be linked with an upregulated EAG (‘linked’ group, **Fig. 4c-d**) while almost two times more elements (2,681, 29%) were located in regions featuring no differential gene expression in primate iAstrocytes (‘not linked’ group, **Fig. 4c-d, Ext. Fig. 5a**). Both the ‘linked’ and ‘not linked’ enhancers were located within the distance of 500 kb of any annotated promoter; elements linked with gene activation featured slightly more openness and a moderately higher level of H3K27ac than elements not linked with differential gene expression in the human lineage (**Ext. Fig. 5b,** *P*=2.3×10^-13^ and *P*=8.8×10^-12^ respectively, two-sided t-test). However, these differences were relatively mild and therefore could not account for the observed functional differences between the two groups of elements.

We explored sequence features that may separate enhancers linked with upregulated EAGs from elements that are not related to transcriptional differences between primate iAstrocytes. An overwhelming majority (∼95%) of the human-chimpanzee sequence differences intersecting the predicted TFBS within both classes of enhancers were single base pair mismatches (**Ext. Fig. 5c**, *Fimo*, **Methods**). These evolutionary changes frequently led to gain in TFBS in the human genome (**Ext. Fig. 5d**). Consistent with their overall poorer conservation (**Fig. 4e**, *** *P*<10^-5^, two-sided t-test), enhancers linked with activation featured significantly more changes in TFBS motifs than elements not linked with DEGs or enhancers active in all primate iAstrocytes (**Ext.Fig. 5c,** two-sided t-test *P*<0.01). Thus, human-specific enhancers linked with activation of gene expression feature signs of stronger positive selection as compared to elements not associated with EAGs or conserved enhancers.

Most enhancers featured at least one sequence change intersecting a prediceted TFBS. To address the contributon of individual TFs to evolution of linked and non-linked enhancers, we compared the frequency of inter-species alterations of the sites recognized by TF located within both classes of enhancers. While for an overwhelming majority of factors, the frequency of TFBS sequence changes was similar for both groups of enhancers (**Fig. 4f**, beige), there was a distinct group of TFs that changed in evolution more frequently when embedded within enhancers linked with upregulated EAGs (blue, **Fig. 4f**, *N*=89, Fisher’s exact test, *P*-adj.<0.01). We repeated the same analysis considering TAD annotation instead of genomic distance (linked enhancers were human-specific enhancers in the same TAD as the up-regulated EAG while not-linked were enhancers gained in humans in TADs that did not feature any evolutionary modulated genes) and obtained analogous result (not shown).

Strikingly, 77 out of 89 motifs changing more frequently at enhancers linked with evolutionary gene upregulation (**Fig. 4f,** P-adj.<0.01), corresponded to cognate sites of the recently described ‘stripe’ TFs^64^, including MAZ, SP1-4, PATZ1 (87.5%, odds=15.3, Fisher’s test *P*<2.2×10^-16^, **Fig. 4g**). These broadly expressed TFs favour binding of other transcriptional regulators and act to ensure proper interaction dynamics of tissue-specific^64^ and housekeeping TFs^65^ with chromatin. Over 77% (1,118) of 1,443 enhancers linked with upregulated EAGs, intersected an evolutionary alteration in motif of at least one ‘stripe’ TF identified in this comparison.

While we observed concomitant evolutionary changes in motifs of TFs other than ‘stripe’ factors within almost all the linked enhancers (1,424 of 1,449 linked enhancers featured such a sequence change), there was no other evident group of TFs that would be hit by an evolutionary change more frequently at linked than the non-linked enhancers. Likewise, evolutionarily lost enhancers linked with gene downregulation often featured an interspecies change in one of the 77 ‘stripe’ TFs identified through the analysis of the gained enhancers (**Ext. Fig. 5e**).

Consistent with our previous analysis (**Ext. Fig. 5d**), the evolutionary changes in motifs recognised by ‘stripe’ TFs were by and large leading to a gain of TFBS in the human genome (**Ext. Fig. 5f)**. Conforming to this observation, comparing the human and chimpanzee versions of the cognate motifs of ‘stripe’ TFs within the human-specific enhancers linked with upregulated EAGs, we found a significant increase in the footprints in the human lineage (**Ext. Fig. 5g**, *HINT* tool, **Methods**). Hence, the evolutionary changes in the DNA sequence of the cognate sites of ‘stripe’ TFs lead to increased binding of these TFs at the human-specific enhancers. Remarkably, virtually all the upregulated EAGs featuring a human-specific enhancer in their vicinity, could be linked with at least one enhancer displaying a change in ‘stripe’ TFBS (**Fig. 4h**) further sustaining the view that changes in ‘stripe’ TFBS contribute to enhancer activity. Altogether, changes in DNA motifs of ‘stripe’ TFs segregate with enhancers linked with positive transcriptional change in evolution.

### Deep learning models the functional impact of evolutionary changes in enhancer sequence

Deep learning methods have been recently deployed to predict the contribution of evolutionary changes in DNA sequence to enhancer activity^66,67^. Hence, to assess the impact of the human-chimpanzee sequence changes in motifs recognised by the 77 ‘stripe’ TFs on enhancer activity, we employed a convolutional neural network (CNN) based on the Basset architecture^68^. The CNN was initially trained on an independent dataset^69^. We re-trained it using the human iAstrocyte regulome, which allowed us to reach 90% prediction power (**Fig. 5a**, **Ext. Fig. 6a**, **Methods**). As expected, human sequences of enhancers that gained openness generated higher CNN response than their chimpanzee counterparts. The inverse was evident at enhancers losing openness (**Fig. 5b**) highlighting the power of our CNN to detect the impact of evolutionary differences in sequence on enhancer activity. Critically, restoration of the chimpanzee version of the ‘stripe’ TF motifs within enhancers linked with gene upregulation led to a significant loss of CNN-predicted activity of these elements (**Fig. 5b**). These data are consistent with the notion that the evolutionary gain in binding sites of ‘stripe’ TFs renders enhancers more active (**Fig. 5c**).

**Figure 5:**
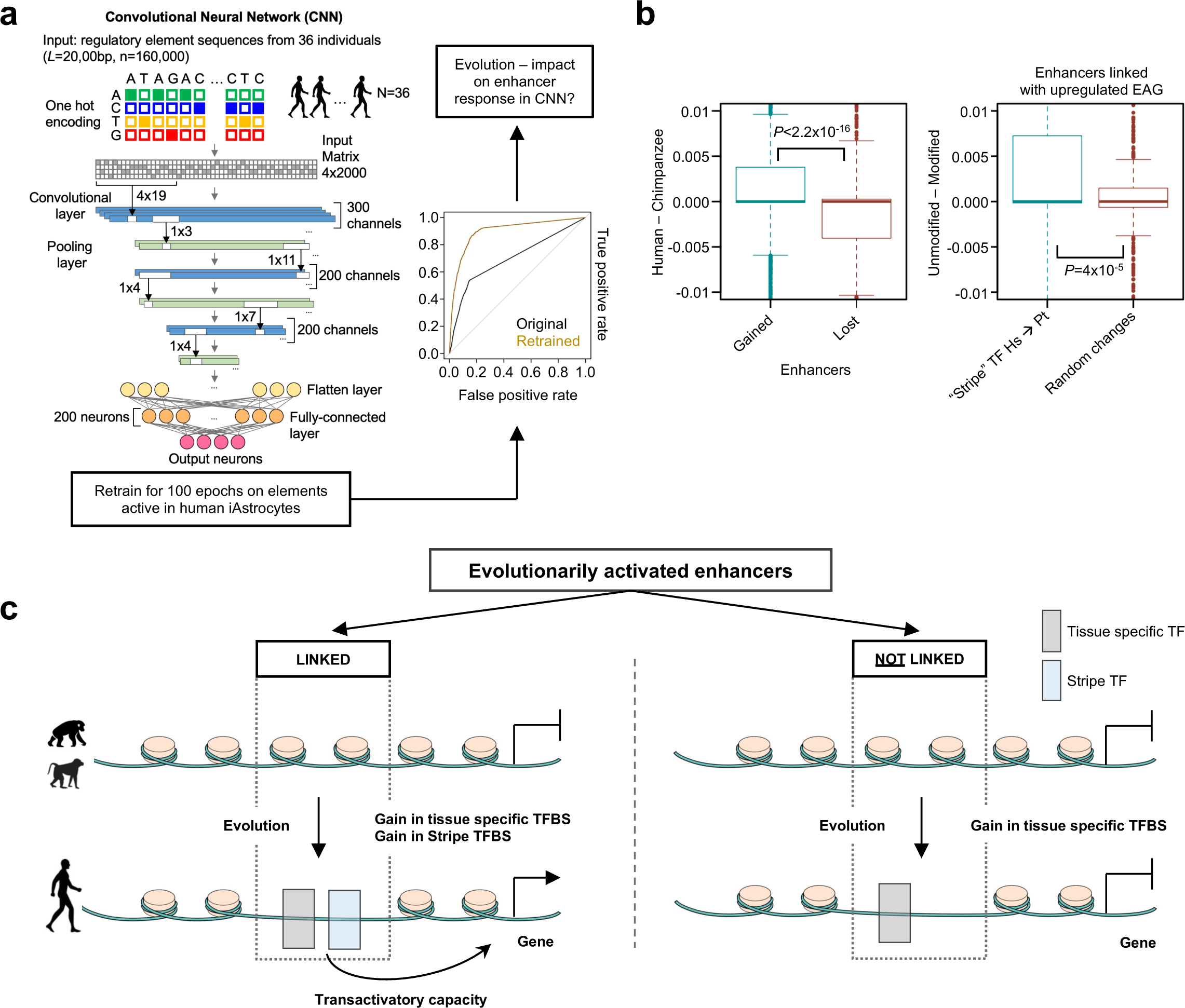
Convolutional neural network predicts the role of stripe transcription factors in evolutionary gain of transactivatory potential of enhancers. **a.** The Convolutional Neural Network (CNN) trained previously on 160,000 regulatory sequences obtained from glioma samples from 36 individuals (69) was considered and retrained for 100 epochs using sequences of elements identified in human iAstrocytes. AUC of the retrained network is increased through re-training. **b.** Human version of the motifs bound by the ‘stripe’ TFs confers a higher regulatory element potential than the chimpanzee version of the same sequence. Left panel: boxplot displaying the change in the CNN response of human and chimpanzee versions of enhancers gained or lost in evolution. Right panel: enhancers linked with activation were considered; boxplot displays the change in CNN response of enhancers induced by the exchange of human motifs bound by ‘stripe’ TFs to the chimpanzee version of the motif. Control group consisted of enhancers linked with upregulated EAGs where sequences were mutated randomly (keeping the total number of mutations equal to the one observed in human-chimpanzee comparison). **c.** Emergence of stripe transcription factor binding sites contributes to the evolutionary gain in transactivatory potential of enhancers.

Altogether, enhancers linked with gain in gene expression feature broad activation of motifs recognised by ‘stripe’ TFs suggesting an underlying role of these widely expressed factors in shaping enhancer activity and thereby gene expression in evolution.

## Discussion

The evolution of the nervous system is characterised by the early segregation of function between neurons, responsible for long-range signaling, and neuroglia performing all defensive and homeostatic tasks^1,70^. In vertebrates, the homeostasis of the nervous system is maintained by astroglia, of which astrocytes are the largest, but not the only representative. The human brain is characterised by highly complex astrocytes^12^, tightly involved in maintenance and regulation of synaptic transmission and hence behaviour^2^, whereas astrocytic malfunction contributes to the pathogenesis of brain diseases^71^. We demonstrate that iPS-derived astrocytes (iAstrocytes) recapitulate the evolutionary advance of morphology in hominids, thereby establishing them as a model to study the evolution of this cell type. The iPS-based systems allow us to unveil the interspecies differences in the regulomes and gene expression profiles, hence revealing mechanisms of evolution of the primate brain^13,72–75^. Our results show the genetic encoding of the differences in the cellular architecture and functional versatility of astrocytes in primates. Dissecting the role of identified genes will help to reveal the molecular pathways of the evolutionary gain in astrocyte cell size and complexity.

Heterozygous deleterious mutations in genes related to chromatin biology or biochemical pathways essential for functions of the nervous system frequently lead to intellectual disability (ID)^76–78^. We find that ID-related genes are transcriptionally downregulated in primate brain evolution. The nature of the trade-off elicited by this process remains largely unclear. Dampened expression of ID-related genes may favour enhanced complexity of the human transcriptome. For instance, the diminished expression of negative regulators of splicing could lead to a more intricate universe of mRNAs produced by the cell. Yet, the transcriptional downregulation of ID-related genes may render humans particularly vulnerable to deficits in their dosage. Addressing the interplay between the level of expression of ID-related genes and the development of the human brain will be instrumental in determining the consequences of the evolutionary loss of transcriptional activity of these loci.

How enhancers gain their *cis*-regulatory potential in evolution is one of the fundamental questions in the field. Mutations of binding sites recognised by specific TFs were previously linked with evolution^61^. However, the existence of more general mechanisms underlying the gain of transactivatory potential of newly evolved enhancers remained unknown. By contrasting evolutionary changes at enhancers that can be linked to transcriptional upregulation, with alterations at elements that are not related to changes in gene expression, we provide evidence that the gain of binding sites of broadly expressed ‘stripe’ transcription factors favour gene activation. Binding of ‘stripe’ TFs is central to stabilise the interactions of lineage-specific TFs with DNA^64^. Therefore, we propose that by introducing ‘stripe’ TFBS to the enhancers, evolution renders them apt in their transactivatory role.

Altogether, we provide an in vitro model to track gradual changes in gene expression and alterations in DNA regulatory elements that contribute to the evolution of astrocytes. Combining evolutionary sequence analysis and machine learning, we show that gain in the binding sites of ‘stripe’ TFs, a group of housekeeping transcriptional regulators with a previously unappreciated role in evolution, is instrumental for eliciting enhancer activity.

## Acknowledgments

Work in the Pękowska lab is funded by the Dioscuri Grant (Dioscuri is a programme initiated by the Max Planck Society (MPG), jointly managed with the National Science Centre in Poland (NCN), and mutually funded by the Polish Ministry of Education and Science and the German Federal Ministry of Education and Research (BMBF)), by the Confocal imaging was performed at the Laboratory of Imaging Tissue Structure and Function which serves as an imaging core facility at the Nencki Institute of Experimental Biology and is part of the infrastructure of the Polish Euro-BioImaging Node. We thank Fred Gage from The Salk Institute for Biological Studies for providing iPS lines. We would like to thank Bożena Kamińska-Kaczmarek and Bartosz Wojtas from the Nencki Insitute. We would like to acknowledge the Municipal Zoological Garden in Warsaw and in particular Dr Agnieszka Czujkowska and Dr Andrzej Kruszewicz.

## Author contributions

Conceptualization: AlePek,; Methodology: KC, AP, IF, AV, RB, BW, AlePek; Investigation: KC, AP, DC, DB, ED, RB, KS, MCK, MN, WK, AlePek; Visualization: KC, AP, DC, BD, AlePek; Funding acquisition: AlePek; Project administration: AlePek; Supervision: AlePek Writing – original draft: KC, AV, AlePek; Writing – review & editing: KC, AV, AlePek **Competing interests:** Authors declare that they have no competing interests.

## Data and materials availability

Raw sequencing data are available in the ArrayExpress database (http://www.ebi.ac.uk/arrayexpress) under following accession numbers: RNA-seq: **E-MTAB-XXXX**; ATAC-seq: **E-MTAB-XXXX**; ChIP-seq H3K27A: **E-MTAB-XXXX**; ChIP-seq H3K4me: **E-MTAB-XXXX**; ChIP-seq CTCF: **E-MTAB-XXXX**; Hi-C: **E-MTAB-XXXX**. Code is available as Supplementary Information.

---

**Extended Figure 1:**
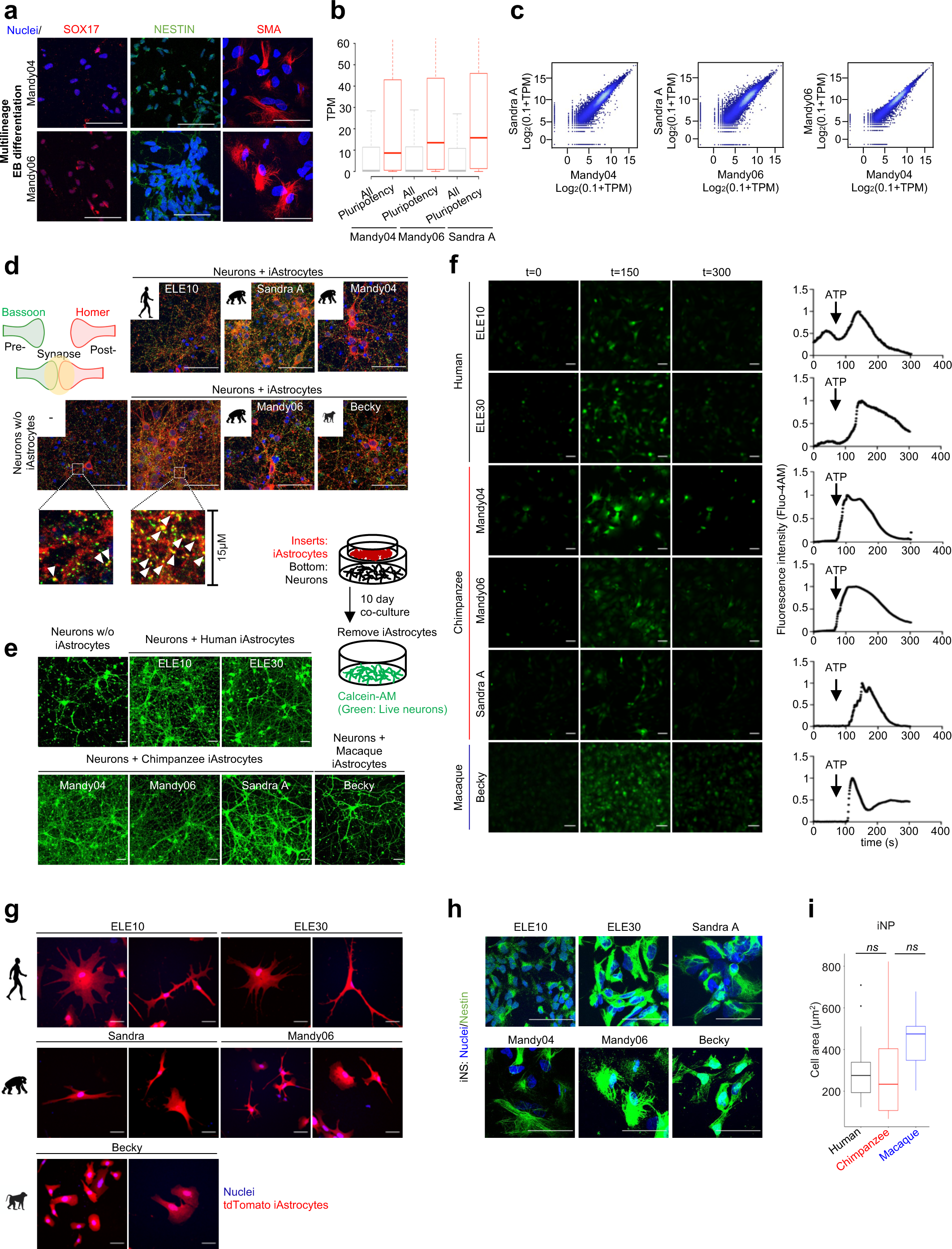
Refers to: Primate iAstrocytes recapitulate evolutionary features of foetal brain astrocytes. **a.** Trilineage differentiation to assess pluripotency of the chimpanzee iPS cell lines Mandy04 and Mandy06 generated in this study. Representative immunofluorescence staining to assess the presence of endoderm marker SOX17 (red); ectoderm marker Nestin (green) and mesoderm marker SMA (red). Both lines differentiate readily to endo-, ecto-, and mesoderm. Scale bar: 50μm. **b.** Expression (transcripts per million; TPM) of pluripotency-related genes and all genes in Mandy04, Mandy06 iPS lines and in a previously validated chimpanzee iPS line SandraA. **c.** Comparison of transcriptomes in chimpanzee iPS cells. Our newly generated lines Mandy04 and Mandy06 feature similar gene expression patten to the established chimpanzee iPS line SandraA. **d.** iAstrocytes promote synapse formation by rat embryonic (E18) neurons. Acutely purified rat embryonic neurons were co-cultured in the presence or in the absence of primate iAstrocytes for 10 days (Methods). Presence of pre-(Bassoon, green) and post-synaptic (Homer, red, inlet cartoon) markers was assessed in the cultures using immunofluorescence. We observed a marked gain in yellow puncta corresponding to synapses. Scale bar in images: 50μm; scale bar in inlets: 10μm. **e.** Primate iAstrocytes promote the survival of rat embryonic neurons (Methods). Calcein (green) staining represents live neurons. Scale bar: 50μm. **f.** iAstrocytes respond to extracellular ATP by generating calcium (Ca2+) signals; Left panel: snapshots of fluorescence signal at 0, 150, 300 seconds of image recording. Scale bar: 50μm; Right panel: average profiles of fluorescence intensity over time (n=50 cells/line). **g.** Human iAstrocytes are bigger than chimpanzee iAstrocytes. Representative examples of iAstrocytes transduced with a vector encoding cytoplasmic tdTomato used for cell area measurement. Scale bar: 50μm. **h.** Primate iNPs do not differ in size. Examples of nestin-stained human, chimpanzee, and macaque iNPs cells used for cell area measurements. Scale bar: 50μm. **i.** Quantification of the cell area of nestin-stained iNP cells (n= 39 (human); 28 (chimpanzee); 11 (macaque), ns).

**Extended Figure 2:**
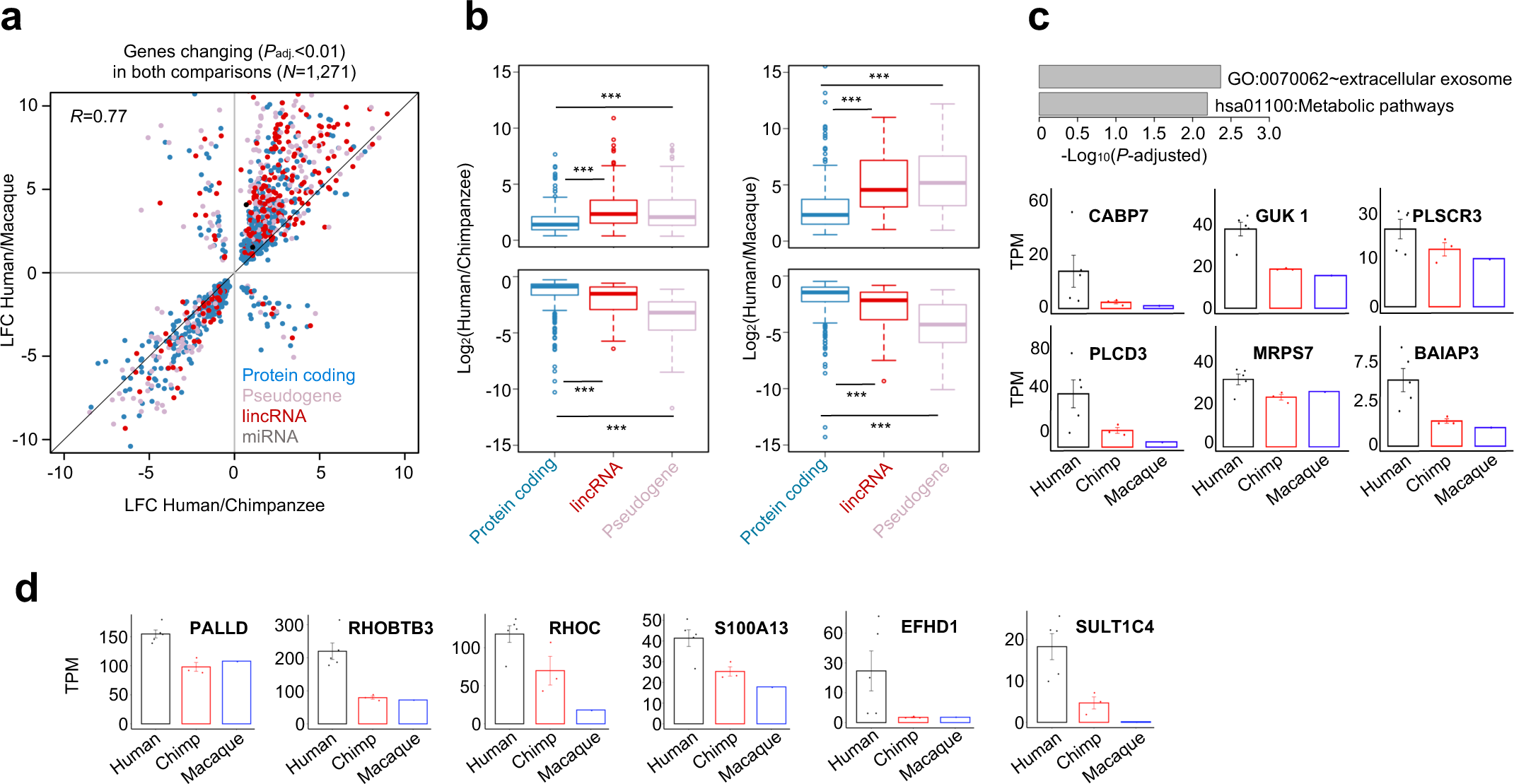
Refers to: Stepwise evolutionary changes in gene expression during astrocyte evolution. **a.** Congruent changes in inter-species comparisons of gene expression in primate iAstrocytes. Considering genes identified as significantly altered in both interspecies comparisons of iAstrocyte transcriptomes (human vs chimpanzee and human vs macaque, differentially expressed genes DEG, *P*-adj.<0.01), the plot displays the comparison of logarithm base 2 of fold changes (LFC) scored when comparing human to chimpanzee (x-axis) and human to macaque (x-axis). **b.** Stepwise transcriptional changes in evolution. Left panel: LFC of gene expression for up- and down regulated EAGs in human versus chimpanzee, right panel and LFC of expression of up- and down regulated EAGs in human vs macaque (right panel). Expression of protein coding genes is less affected in evolution than the expression of non-protein coding loci (*** *P*-val.<10^-10^, two-sided t-test). The LFC of gene expression is significantly stronger when comparing human to macaque iAstrocytes than when comparing human to chimpanzee iAstrocytes (*P*-val.<2.2×10^-16^, two-sided t-test not displayed in the figure, compare corresponding boxplots in left and right panels). **c.** Human astrocytes feature frequent upregulation of genes related to metabolism and extracellular exosome. Gene Ontologies and KEGG pathways enriched in the list of upregulated EAGs (Benjamini Hochberg corrected *P*-values, https://david.ncifcrf.gov/). Bottom: expression of example EAGs related to metabolism in primate iAstrocytes **d.** Normalized expression of exemplary EAGs related to cytoskeleton and Ca^2+^ signalling in primate iAstrocytes.

**Extended Figure 3:**
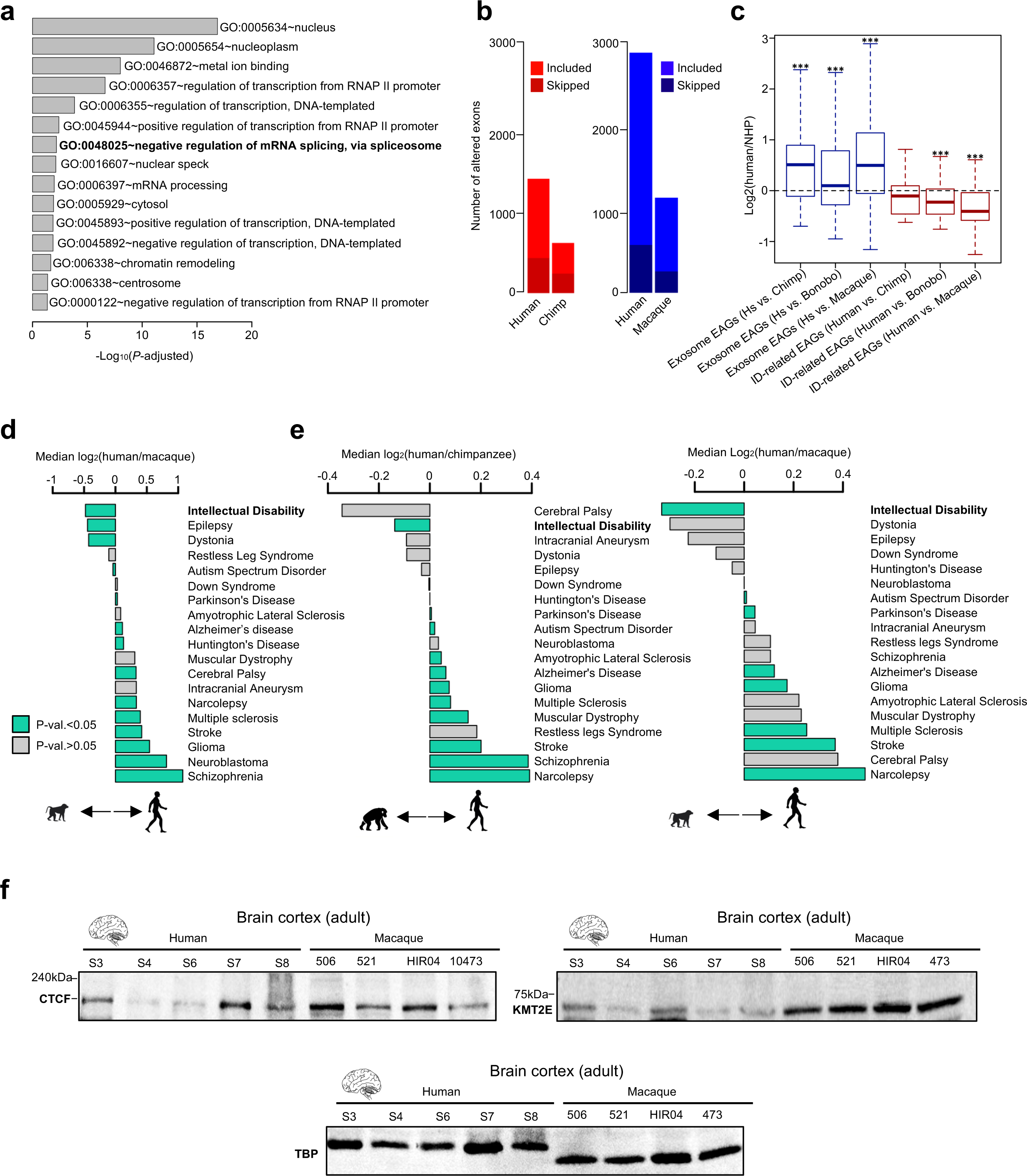
Refers to: Intellectual disability gene s are frequently transcriptionally downregulated in the evolution of the human brain. **a.** Gene Ontologies enriched in the list of genes downregulated EAGs obtained through analysis using DAVID tool (https://david.ncifcrf.gov/). **b.** Human iAstrocytes display increased number of splicing events comparing to NHP. Included and skipped exons in iAstrocytes identified in the comparison of human and chimpanzee transcriptomes (left panel, red) and human versus macaque transcriptomes (right panel; blue). **c.** Distribution of fold changes of expression for EAGs related to extracellular exosome and ID (loci identified in Fig. 2 e,h). Previously published dataset corresponding to bulk RNA-seq profiling of human, chimpanzee, gorilla, and macaque post-mortem cortical tissues^21^ was considered in this analysis. **d.** Median LFC of expression of disease-related genes in adult human versus macaque brain cortex. For consistency, we considered only diseases for which there were more than 30 genes linked in the BrainBase database. The ID-related genes (n=138), are significantly downmodulated in adult human vs macaque cortex (turquoise: *P*-val.<0.05; grey: *P*-val.>0.05). **e.** Median LFC of expression of disease-related genes in adult human versus NHP brain cortex. For consistency, we considered only diseases for which there were more than 30 genes linked in the database. Analysis based on bulk RNA-seq profiles (14) of adult female human, chimpanzee, and macaque normal cortical tissue. ID-related genes (n=138, BrainBase), are significantly downregulated in all the comparisons (turquoise: *P*-val.<0.05; grey: *P*-val.>0.05). **f.** Level of CTCF (left panel) and KMT2E proteins in human and macaque cortical tissues. Exemplary Western blot analysis of whole tissue protein extract. TBP was used as loading control (bottom panel).

**Extended Figure 4:**
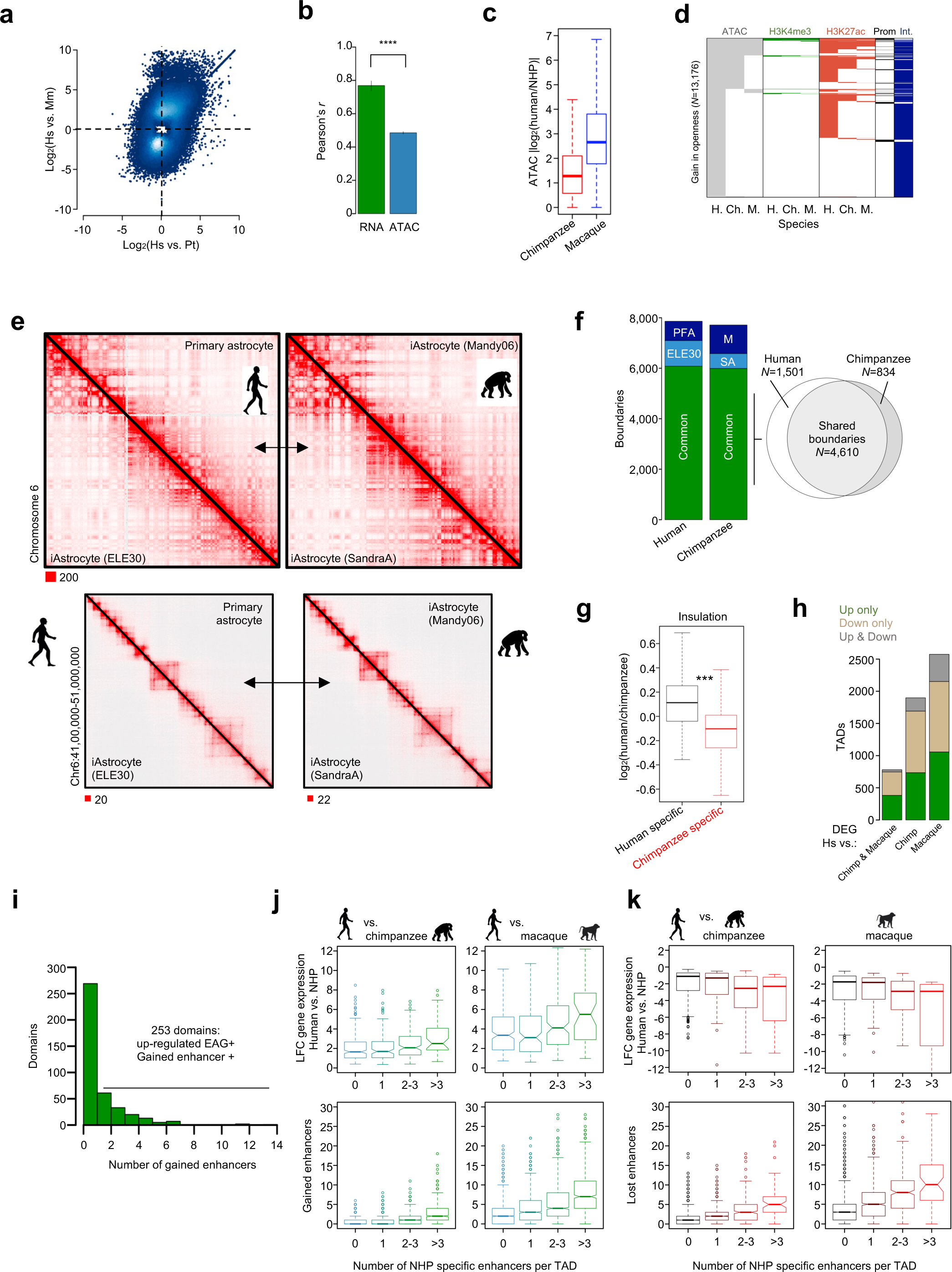
Transcriptional and regulatory changes within topologically associating domains in primate astrocyte evolution. **a.** Correlation of changes in chromatin openness between human and chimpanzee and human and macaque iAstrocytes. LFC of normalized ATAC-seq signal at DORs identified in both inter-species comparison (*P*-adj.<0.1). **b.** Changes in transcriptome between human and chimpanzee and human and macaque are more correlated with each other than are the between-species changes in chromatin openness (P-value<2.2×10^-16^, two-sided t-test). **c.** Stepwise gain of openness at human-specific DORs (*P*-adj.<0.1 in both comparisons). The LFC increases with the more pronounced evolutionary distance between species. **d.** Chromatin and genomic signature of DORs featuring a significant increase in ATAC-seq signal in human (H.) as compared to chimpanzee (Ch.) and macaque (M.) iAstrocytes. Colour indicates overlap with peaks (Gray – ATAC, Green H3K4me3, Red H3K27ac) or genomic annotation (black: overlap with a promoter region; blue: no overlap with promoter region). **e.** *In-situ* Hi-C profiles of chromosome 6 in human and chimpanzee cells (upper panel: entire chromosome 6, lower panel: zoom into a randomly chosen region of chromosome 6 in human and its corresponding region in the chimpanzee). We observe large-scale conservation of chromatin structure between humans and chimpanzees. **f.** TAD boundaries are highly reproducible between human iAstrocytes and primary foetal astrocytes and in the two iAstrocyte lines in chimpanzees and between species. Classification of TAD boundaries common or specific to species. Boundaries were called using TopDom. We considered only boundaries identified in both samples in each species in the Venn diagram. **g.** Insulation strength changes for human- and chimpanzee-specific TAD boundaries. There is a significant increase of insulation in human cells at borders gained in human iAstrocytes and a decrease of IS at borders lost in the human lineage as compared to chimpanzee (\*\*\**P*-val.<2.2×10^-16^, two-sided t-test). **h.** TADs contain genes changing expression level in the same direction in evolution. Displayed is the number of TADs containing only up-or downregulated EAGs and the number of TADs containing both up- and downregulated genes. The three bars correspond to analyses considering EAGs (left) or differentially expressed genes in human versus chimpanzee (middle) or human versus macaques (right bar). **i.** TADs contain genes changing expression level in the same direction in evolution. Displayed is the number of TADs containing only up- or downregulated EAGs and the number of TADs containing both up- and downregulated genes. The three bars correspond to analyses considering EAGs (left) or differentially expressed genes in human versus chimpanzee (middle) or human versus macaques (right bar). **j.** LFC of expression of upregulated EAGs scales with the per-TAD number of enhancers gained in evolution. TADs were stratified by the number of enhancers gained in human as compared to both chimpanzee and macaque cells (x-axis). Top panel: LFC of expression (y-axis, *DESeq2* method) of genes annotated for each group of TADs; human versus chimpanzee (left); human versus macaque (right). Bottom panel: number of enhancers gained in humans as compared chimpanzee (left) or macaque (right). Since TADs were stratified by the number of enhancers gained in the two comparisons; these elements were excluded from analysis presented in the bottom panel. **k.** LFC of expression of downregulated EAGs scales with the per-TAD number of enhancers lost in evolution. TADs were stratified by the number of enhancers lost in the human cells as compared to both chimp and macaque iAstrocytes (x-axis). Top panel: expression LFC (y-axis, *DESeq2* method) of genes, annotated for each group of TADs, in the comparison of human versus chimp (left) or of human versus macaque (right). Bottom panel: number of enhancers lost in humans as compared chimpanzee (left) or macaque (right). Since TADs were stratified by the number of enhancers lost in the two comparisons, these elements were excluded from analysis presented in the bottom panel.

**Extended Figure 5:**
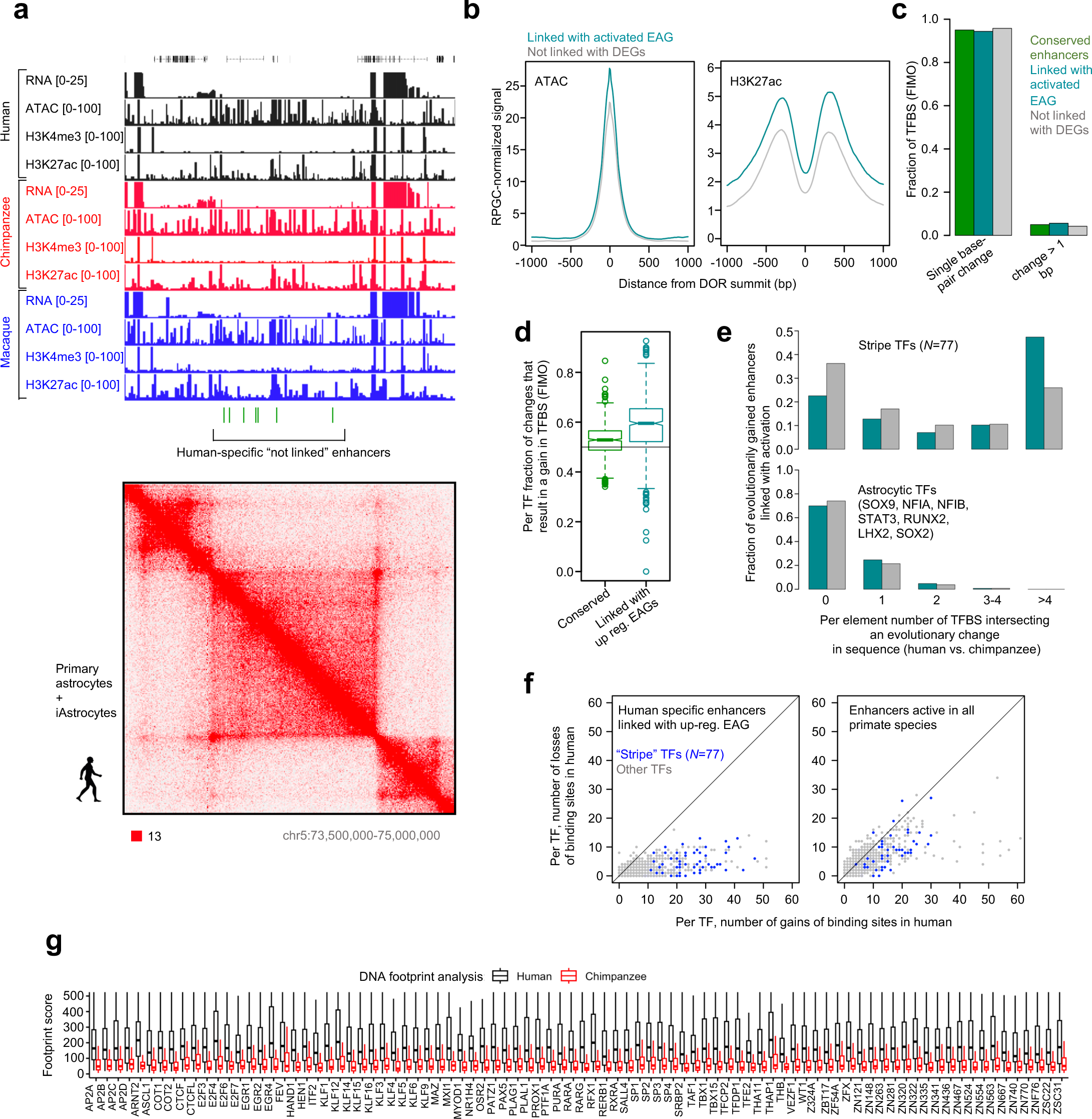
Referring to: Gain of binding sites of ‘stripe’ transcription factors at enhancers linked to gene activation. **a.** Gene expression, chromatin activity and architecture at a (randomly chosen) locus at chromosome 5, where we did not score significant transcriptional changes in primate iAstrocytes despite the presence of a cluster of human specific enhancers. Green bars: enhancers gained in the human iAstrocytes; upper panel: RNA-, ATAC- and ChIP-seq tracks aligned to the consensus genome; lower panel: in-situ-Hi-C reads aligned to the human genome. **b.** Enhancers linked with gene up-regulation in evolution feature higher openness and a higher level of H3K27ac mark compared to elements not linked with differential gene expression in the primate iAstrocytes. Average profile of ATAC-seq (left) or ChIP-seq for H3K27ac (right) at human specific enhancers linked upregulated EAGs (turquoise) or not linked with activation (grey). **c.** Great majority of evolutionary changes in the DNA sequence at the predicted TFBS are single base pair alterations. **d.** Changes in the sequences of the predicted TFBS lead, by and large, to a gain in TFBS in humans as compared to chimpanzee. TFBS were predicted by FIMO. **e.** Number of enhancers featuring evolutionary changes in TFBS of ‘stripe’ TFs (upper panel) or in the cognate motifs the established astrocytic TFs (lower panel, turquoise: enhancers liked with DEGs; grey: enhancers not linked with DEGs; 77 ‘stripe’ TFs identified in panel d were considered in this analysis). **f.** Evolutionary changes in the DNA sequence of linked enhancers lead to gains rather than losses of TFBS. Displayed is the number of gained sites per TF motif (that is sites called in the human genome and not in the chimpanzee) and lost sites (TFBS called in the chimpanzee but absent in the human genome) for linked enhancers (left panel) and enhancers conserved in evolution (right panel). **g.** ATAC-seq based footprint analysis of ‘stripe’ TFBS affected by evolution and identified within enhancers linked with upregulated EAGs confirms stronger signal in human than in the chimpanzee iAstrocytes. 10 bp intervals centered on the midpoint of the motif annotated in the human were considered for the footprint analysis. These genomic intervals were lifted over to the chimpanzee genome.

**Extended Figure 6:**
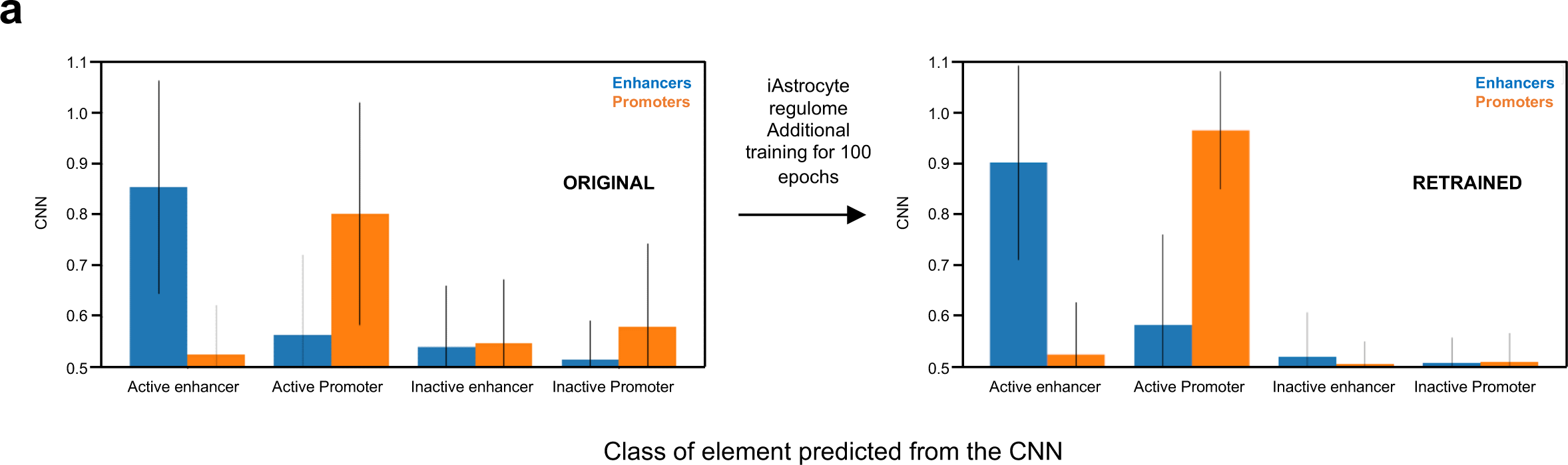
Referring to: Convolutional neural network captures the impact of evolutionary changes in sequence on enhancer activity. **a.** Prediction power of the CNN before and after retraining.

